# Tuning trophoblast invasion in a gelatin hydrogel via soluble cues from the maternal-fetal interface

**DOI:** 10.1101/2020.04.16.045484

**Authors:** Samantha G. Zambuto, Kathryn B.H. Clancy, Brendan A.C. Harley

## Abstract

Trophoblast cells play multiple critical roles in pregnancy, notably modulating blastocyst attachment to the endometrium as well as invading into and actively remodeling the endometrium to facilitate biotransport needs of the growing embryo. Despite the importance of trophoblast invasion for processes essential at early stages of pregnancy, much remains unknown regarding the balance of signaling molecules that may influence trophoblast invasion into the endometrium. The goal of this study was to use a three-dimensional trophoblast spheroid invasion assays to examine the effect of cues from the maternal-fetal interface on trophoblast invasion. We report use of a methacrylamide-functionalized gelatin (GelMA) hydrogel to support quantitative analysis of trophoblast outgrowth area and cell viability. We show this multidimensional model of trophoblast invasion can resolve quantifiable differences in outgrowth area and viability in the presence of a known invasion promoter, epidermal growth factor, and a known invasion inhibitor, transforming growth factor β1. We then investigate the sensitivity of trophoblast invasion to cortisol, a hormone associated with exogenous stressors. Together, this approach provides a toolset to investigate the coordinated action of physiological and pathophysiological processes on early stages of trophoblast invasion.

**IMPACT STATEMENT:** Trophoblast cells from the invading blastocyst play crucial roles in pregnancy, including remodeling endometrial structure to support embryonic biotransport needs; however, much remains unknown regarding the balance of signaling molecules that may influence trophoblast invasion. We show this multidimensional model of invasion can resolve quantifiable differences in outgrowth area and viability in the presence of known invasion promoters and inhibitors and then investigate invasion sensitivity to cortisol, a hormone associated with exogenous stressors. This approach provides a toolset to investigate the coordinated action of molecules from the maternal-fetal interface on trophoblast invasion that may be challenging to study in humans.

## 1. Introduction

Pregnancy is a complex biological process that involves molecular dialogue between trophoblast cells from the invading blastocyst and cells from the target of implantation, the endometrium. This dialogue coordinates the extent of trophoblast invasion into the endometrium. Although variations in trophoblast invasion are believed to impact the success of a pregnancy (1–4), much remains unknown regarding pathological signaling processes that drive trophoblast invasion into the endometrium. Implantation occurs when the blastocyst establishes a stable connection with the endometrium (1, 2). In order for implantation to occur, the endometrium must undergo preparation via hormonal priming and enter what is known as the implantation window, a short 4-day window during the mid-secretory phase of a 28-day menstrual cycle (1, 5). Human implantation is thought to occur in three phases: apposition, adhesion, and invasion (2, 5). Apposition is defined as initial, unstable attachment of the blastocyst to the endometrial luminal epithelium (1, 5). Adhesion initiates physical interactions between trophoblast cells from the blastocyst and endometrial epithelium (1, 5). Finally, invasion occurs when trophoblast cells breach the endometrial epithelium and subsequently invade into the underlying stroma (1, 5). Perturbations in the implantation processes can result in a variety of pregnancy disorders. Implantation failure accounts for approximately 75% of failed pregnancies and represents a significant challenge to fertility (1, 2). Implantation failure is not clinically recognized as a pregnancy and defective implantation likely causes adverse effects that compound over the course of the pregnancy. These defects can result in poor pregnancy outcomes, including the hypertensive pregnancy disorder preeclampsia, intrauterine growth restriction, and recurrent pregnancy loss (2). Because implantation involves a highly coordinated molecular dialogue between endometrial cells and trophoblast cells, developing a deeper understanding of the biological mechanisms surrounding implantation may provide critical insights into pregnancy and pregnancy disorders.

Implantation has never been observed in humans due to ethical concerns regarding studying pregnancy in humans as well as a lack of tools to study this process in the body (1, 6, 7). The blastocyst is fully embedded in the endometrial stroma by approximately 10 days post-conception which provides a unique challenge to obtaining direct mechanistic evidence regarding what influences trophoblast invasion into the endometrium (1). Rare histological specimens have allowed us to glean some information on implantation in human specimens; however, only a limited number of samples exist and these specimens cannot provide information on implantation in real time (1, 6). Additionally, inferring mechanistic processes from animal models may not be accurate due to significant differences between human and animal pregnancy, even amongst humans and non-human primates (7). Although we can use two-dimensional assays, including wound healing assays and Boyden chamber assays, to probe biological mechanisms of implantation, these traditional types of invasion assays cannot recapitulate the complexity necessary to capture dynamic processes associated with trophoblast invasion such as matrix remodeling. Tissue engineered models allow for mechanistic studies using rare populations of cells cultured for extended periods of time in three-dimensional environments. Such models can replicate relevant biophysical properties inspired by the native tissue, including matrix stiffness, extracellular matrix composition, and three-dimensional architecture. Advanced tissue engineered platforms have increasingly been utilized to study trophoblast invasion and migration in three-dimensional biomaterial platforms (8–11); however, some limitations still remain with existing models. The use of three-dimensional bioprinting (8, 9) and microfluidic technology (10) allows for tracking migration of trophoblast cells toward soluble factor gradients within biomaterials but these models quantify migration in dissociated cells and hence, do not recapitulate the native spherical structure of an invading blastocyst. Existing models that utilize embryos (11) utilize Matrigel which contains a heterogeneous combination of extracellular matrix proteins and exhibits significant lot to lot variability (12, 13). Further, these models did not quantify trophoblast invasion from the embryo. Adaptation of tissue engineered platforms of increasing complexity can address these issues by employing homogeneous, well-characterized materials and replicating native tissue structure and they can provide additional tools to study core processes associated with trophoblast invasion as they relate to pregnancy and pregnancy disorders.

The overall objective of this study was to use methacrylamide-functionalized gelatin (GelMA) hydrogels and advanced trophoblast spheroid invasion assays to quantify trophoblast invasion and cell viability in the presence of cues from the maternal-fetal interface. We show this multidimensional model can quantify trophoblast spheroid outgrowth area and viability using known promoters (epidermal growth factor; EGF) and inhibitors (transforming growth factor β1; TGFβ1) of trophoblast invasion to demonstrate the relevance of our platform for such studies. EGF is a decidual factor which has been shown to stimulate trophoblast migration (8, 14, 15). Transforming growth factor β superfamily members are expressed in the endometrium, with TGFβ1 present in endometrial epithelial and stromal cells (16). Production and secretion of TGFβ by epithelial cells during the secretory phase suggest that it may play a role in implantation (16). Next, we investigate the effects of cortisol, a steroid hormone produced in response to stress, on trophoblast invasion and quantify trophoblast spheroid outgrowth area and viability. Cortisol is a steroid hormone often used to assess the stress response and is relevant to a variety of processes of pregnancy (17, 18). Stressors during pregnancy, including poverty, intimate partner violence, lack of social support, and racism, have been associated with increased risk for certain pregnancy disorders, such as preterm birth, low birth weight, and preeclampsia (19–22). Nonetheless, poorly understood processes of how stress affects early events in pregnancy such as trophoblast invasion motivate our use of a tissue engineered platform to investigate the role of soluble cues from the maternal-fetal interface on trophoblast invasion.

## 2. Materials and Methods

### 2.1. Hydrogel Fabrication and Characterization

#### 2.1.1 Methacrylamide-Functionalized Gelatin (GelMA) Synthesis and Hydrogel Fabrication

Methacrylamide-functionalized gelatin with 57% degree of functionalization, determined by ^1^H-NMR, was synthesized as described previously using the one-pot method developed by Shirahama et al. (23, 24). Lyophilized GelMA (23) was dissolved in phosphate buffered saline (PBS; Lonza, 17-516F) at 37°C to make 5 wt% polymer solutions. 0.1% w/v lithium acylphosphinate (LAP) was used as a photoinitiator (25). Unless otherwise noted, 20 μL prepolymer solution was pipetted onto custom circular Teflon molds (5 mm diameter, 1 mm height). Hydrogels were polymerized under UV light (λ=365 nm, 7.14 mW cm^−2^; AccuCure Spot System ULM-3-365) for 30 seconds.

#### 2.1.2 Hydrogel Characterization

Hydrogel Young’s modulus was determined via compression testing using an Instron 5943 mechanical tester with a 5N load cell (26). Hydrogel disks (10 mm diameter, 2 mm height, 100 μL prepolymer solution), fabricated to have approximately the same height as those made using the smaller molds, were submerged in PBS and allowed to swell for 2 hours at 37°C. Samples were compressed at a rate of 0.1 mm/min and moduli were quantified from the linear region of the stress-strain curve using a custom MATLAB code that calculates modulus from the linear regime at a load of 0.003 N and offset of 2.5% strain to ensure contact with the hydrogel surface. To calculate mass swelling ratio, hydrogels were fabricated using the smaller molds and were hydrated in PBS overnight at 37°C. Swollen hydrogels were weighed, lyophilized, and weighed once more to determine dry mass. Mass swelling ratio was calculated using the ratio of wet polymer mass to dry polymer mass as previously described (23).

### 2.2. HTR-8/SVneo Spheroid Invasion Assays

#### 2.2.1 HTR-8/SVneo Cell Maintenance

HTR-8/SVneo trophoblast cells (ATCC® CRL-3271, used experimentally before passage 6 after purchase) were maintained as per the manufacturer’s instructions in phenol red-free RPMI-1640 supplemented with 5% charcoal-stripped fetal bovine serum (Sigma-Aldrich, F6765) and 1% penicillin/streptomycin (Thermo Fisher, 15140122). All cultures were grown in 5% CO_2_ incubators at 37°C. Routine mycoplasma testing was performed every 6 months to ensure cell quality using the MycoAlert™ Mycoplasma Detection Kit (Lonza).

#### 2.2.2 Spheroid Invasion Assays

Spheroid invasion assays were performed as previously described by our group (23, 27). 4,000 HTR-8/SVneo cells were added to round bottom plates (Corning, 4515) and placed on a shaker for 48 hours to form spheroids. 4,000 cells/spheroid was selected because this generated spheroids with diameters similar to that of an invading blastocyst which ranges from approximately 100 – 200 μm (Supplemental Information Figure 2) (28). Individual spheroids were pipetted onto the Teflon hydrogel molds. Prepolymer solution was added to the mold and spheroids were gently moved to the center of the mold using a pipette tip. Hydrogels were then polymerized and added to 48 well plates containing 500 μL of cell medium per well. Once all spheroids were encapsulated, spheroids were imaged and medium was replaced with 800 μL of medium with or without (control) biomolecules per well. Medium was not changed at any other times during the experiment unless otherwise noted. Spheroids and encapsulated spheroids were cultured in phenol red-free RPMI-1640 supplemented with 2% charcoal-stripped fetal bovine serum, 1% penicillin/streptomycin, and relevant biomolecules if applicable. No differences in cell growth or morphology were found for cells cultured in medium with 2% fetal bovine serum (Supplemental Information Figure 1). Recombinant human transforming growth factor β1 (TGFβ1; R&D Systems, 240-B) and recombinant human epidermal growth factor (EGF; Sigma-Aldrich, E9644) were added to the medium at a concentration of 5 ng/mL. Cortisol (Sigma-Aldrich, H0888) was added to the medium at concentrations of 5, 20, 75, and 150 ng/mL. Control samples were incubated with no added biomolecules. Spheroids were imaged daily using a Leica DMI 4000 B microscope (Leica Microsystems). Total outgrowth area was calculated using the Measure tool in Fiji. Spheroids were manually traced three times and outgrowth area was determined from the average of these three measurements.

**Figure 1.**
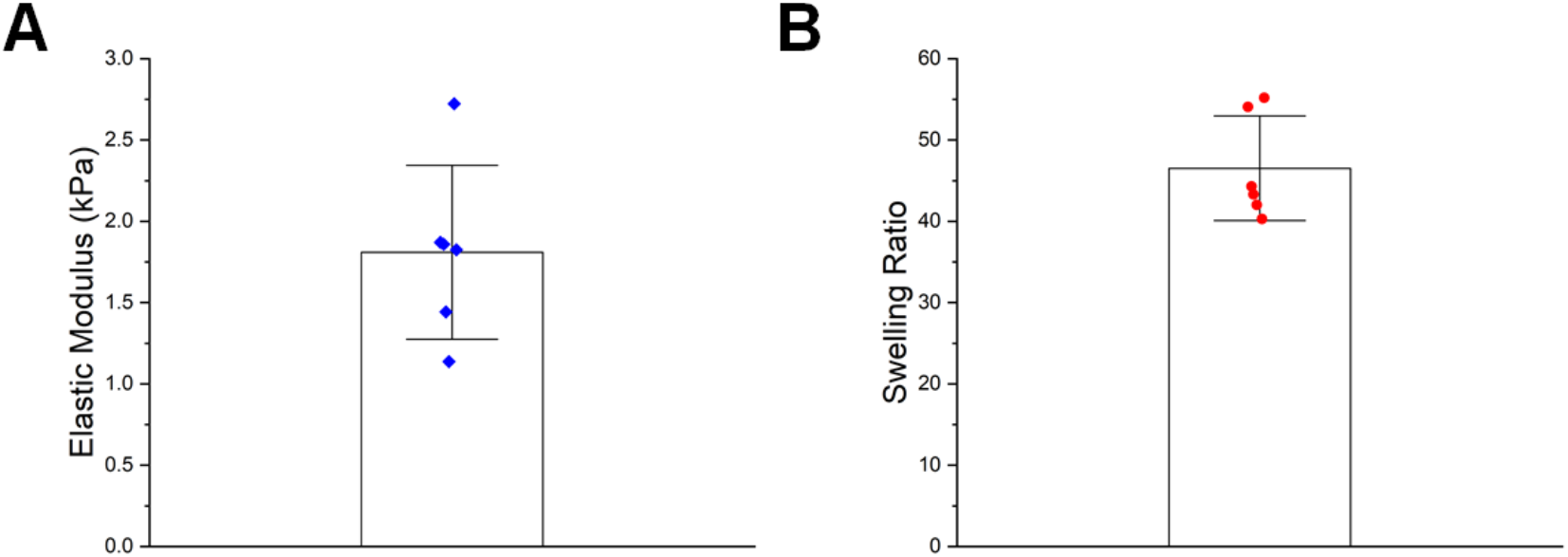
Mechanical characterization of 5 wt% GelMA hydrogels. A) Elastic modulus determined by compression testing (n=6). B) Mass swelling ratio determined by comparing swollen polymer mass to dry polymer mass (n=6). Data presented as mean ± standard deviation with individual data points shown.

#### 2.2.3 Viability Assay

The CellTiter-Glo® 3D Cell Viability Assay (Promega) was used to quantify spheroid viability on day 3. Samples were equilibrated to room temperature for at least 30 minutes prior to running the assay. A stock solution of 1:1 cell medium and CellTiter-Glo® was prepared, medium was removed from each sample well, and 400 μL of the stock solution was added to each sample. Samples were protected from light and incubated for 1 hour on a shaker at room temperature. 100 μL triplicates were added to an opaque plate and luminescence was read immediately using a plate reader (BioTek Synergy HT Plate Reader and Gen5 software; BioTek Instruments, Inc.). A blank was prepared using stock solution. Relative luminescence for each sample was calculated by subtracting the average luminescence value from the blank wells from the average luminescence values of each sample.

### 2.3. Imaging Techniques

#### 2.3.1 Spheroid Staining

Spheroids were encapsulated in hydrogels and grown for 1 or 3 days. Samples were fixed in 4% formaldehyde in PBS for 15 minutes followed by 3 PBS washes. The following solutions were prepared: Permeabilizing Solution (0.1% Tween 20 (Fisher Scientific, BP337) in PBS) and Working Solution (1 μL Phalloidin-iFluor 488 Reagent (Abcam, ab176753) per 1 mL 1% bovine serum albumin (Sigma-Aldrich, A4503) in PBS). All subsequent steps were performed at room temperature on a shaker. Samples were permeabilized in Permeabilizing Solution for 15 minutes. After permeabilization, 300 μL of Working Solution was added per well and samples were protected from light and incubated for 90 minutes. Staining was followed by 4×20 minute PBS washes. Samples were then stained with Hoechst (1:2000 in PBS; Thermo Fisher, H3570) for 30 minutes followed by a PBS wash. Samples were stored in PBS at 4°C until imaged. One Z-stack per spheroid was taken using a Zeiss LSM 710 Confocal Microscope. Maximum intensity projection images were generated using ZEN (blue edition; Zeiss) and Median filtering was used to smoothen images.

#### 2.3.2 Two-Dimensional Immunofluorescent Staining

To determine glucocorticoid receptor expression in HTR-8/SVneo cells, HTR-8/SVneo cells were seeded on 6-well plates at a density of 3 × 10^4^ cells/cm^2^ and cultured in cell growth medium supplemented with 2% charcoal-stripped fetal bovine serum and 1% penicillin/streptomycin until approximately 80% confluence. Cells were fixed using 4% formaldehyde for 15 minutes followed by three PBS washes. Cells were permeabilized in 0.5% Tween 20 in PBS for 15 minutes followed by 3×5 minute washes in 0.1% Tween 20 solution in PBS, blocked in blocking solution (2% bovine serum albumin and 0.1% Tween 20 solution in PBS) for 1 hour at room temperature, and incubated in primary antibody solution (Abcam ab3578 rabbit polyclonal anti-glucocorticoid receptor antibody; 1:20) diluted in blocking solution overnight at 4°C. Following the overnight incubation, 5×5 minute washes in 0.1% Tween 20 solution in PBS were performed and samples were incubated in secondary antibody solution (Thermo Fisher A-21428 Goat Anti-Rabbit IgG (H+L) Cross-Adsorbed Secondary Antibody AlexaFluor 555; 1:500) diluted in blocking solution overnight 4°C while protected from light. 5×5 minute washes in 0.1% Tween 20 solution in PBS were performed followed by a 30-minute stain with Phalloidin-iFluor 488 Reagent diluted in Working Solution. 3×5 minute PBS washes were performed followed by a 10-minute Hoechst (1:2000) stain and a quick wash with 0.1% Tween 20 solution in PBS. Samples were stored in 0.1% Tween 20 solution in PBS at 4°C until imaged. Two images per well (n=3 wells each condition) were imaged using identical image settings on a DMi8 Yokogawa W1 spinning disk confocal microscope outfitted with a Hamamatsu EM-CCD digital camera (Leica Microsystems). Images were pseudo-colored and overlaid using Fiji.

### 2.4. Statistics

OriginPro 2019 (Origin Lab) and RStudio were used for statistical analysis. Normality was determined using the Shapiro-Wilkes test and homoscedasticity (equality of variance) was determined using Levene’s test. For all quantitative data, n=4−6 spheroids were analyzed per sample group. For each experiment, outgrowth area was compared between groups on the same day. Normal, homoscedastic data was analyzed using a one-way ANOVA followed by post hoc Tukey Test. For data that violated the assumption of normality but maintained homoscedasticity, the Kruskal-Wallis Test was used to analyze data followed by Dunn’s post hoc test. For normal, heteroscedastic data, Welch’s ANOVA was used to analyze data followed by the Games-Howell post hoc test. For non-normal, heteroscedastic data, Welch’s Heteroscedastic F Test with Trimmed Means and Winsorized Variances was used to analyze data followed by Games-Howell post hoc test. Significance was set as *p*<0.05. Outgrowth area results are reported as mean ± standard deviation and cell viability data are reported as box plots.

## 3. Results

### 3.1. The development of three-dimensional trophoblast spheroid invasion assays allows for quantitative invasion analysis

We cultured HTR-8/SVneo trophoblast spheroids in hydrogels with mechanical properties (Elastic modulus: 1.8±0.5 kPa; Mass Swelling Ratio: 46.5±6.4) similar to that of native tissue (Fig. 1), which was found to be approximately 1.25 kPa for the decidua basalis and 0.171 kPa for the decidua parietalis (29). We cultured spheroids for up to 3 days and observed cell invasion into the surrounding hydrogel matrix over time (Fig. 2). These assays provide a platform to quantitatively analyze trophoblast invasion in three-dimensions. We subsequently examined differences in invasion and viability in response to exogenous soluble factors.

**Figure 2.**
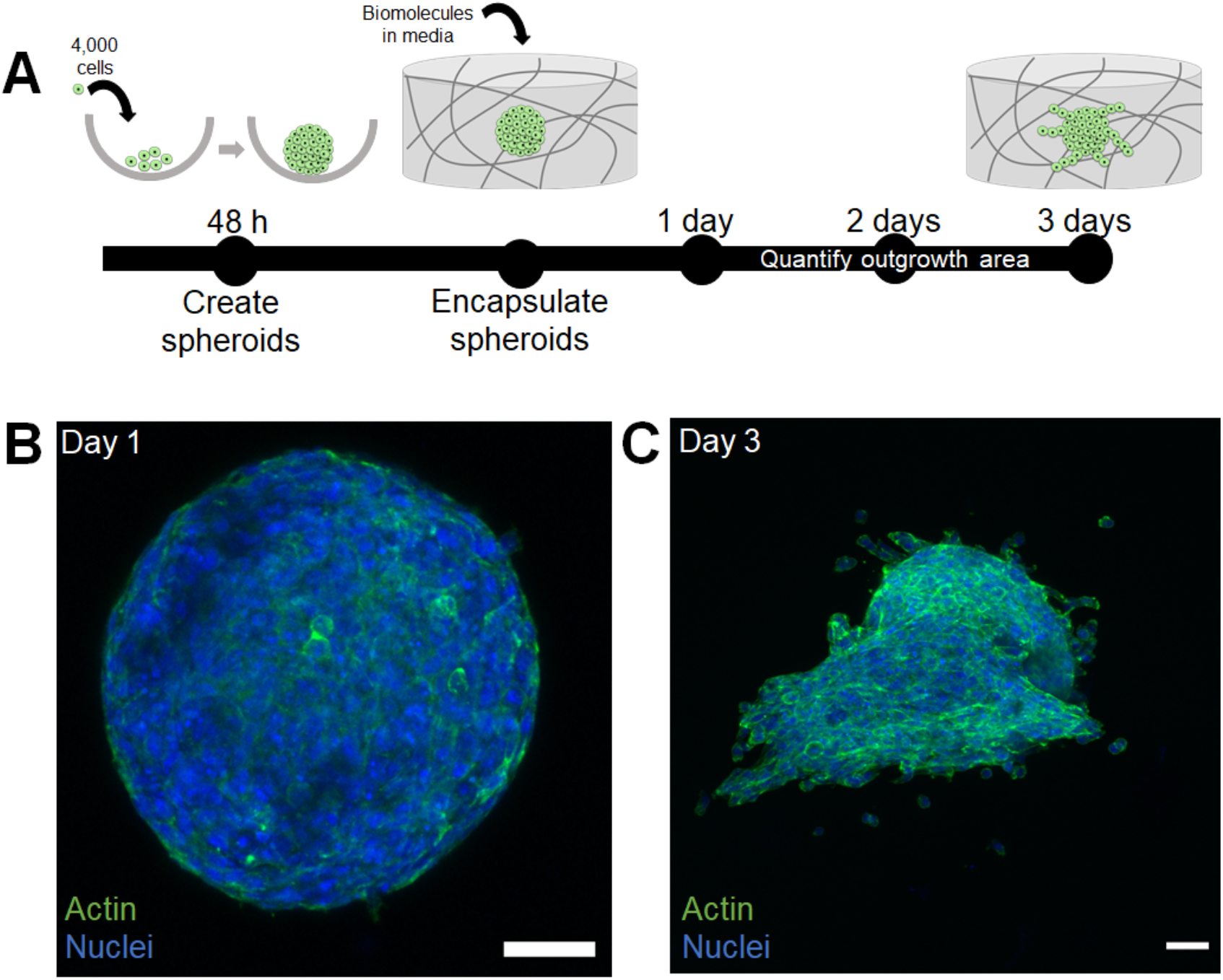
Three-dimensional trophoblast invasion assays in methacrylamide-functionalized gelatin hydrogels. A) HTR-8/SVneo trophoblast cells were seeded into round bottom plates for 48 hours to create spheroids. Spheroids were then encapsulated in hydrogels and allowed to invade the surrounding hydrogel for up to 3 days in the presence of absence of soluble biomolecules in growth medium. B) Representative maximum intensity projection image of a spheroid in a hydrogel 1 day post-embedding. C) Representative maximum intensity projection image of a spheroid 3 days post-embedding. Green: Phalloidin (Actin). Blue: Hoechst (Nuclei). Scale: 50 μm.

### 3.2. Epidermal growth factor and transforming growth factor β1 modulate trophoblast invasion and viability

We quantified shifts in trophoblast invasion and viability via the addition of soluble biomolecules in cell media. Trophoblast spheroids showed increased invasion and increased viability in the presence of known trophoblast invasion promoter epidermal growth factor (EGF). Trophoblast spheroids showed inhibited invasion and decreased viability in the presence of known trophoblast invasion inhibitor transforming growth factor β1 (TGFβ1) by day 2 (Fig. 3). Starting by day 1, the TGFβ1 condition was significantly different compared to control (Welch’s ANOVA: *p*=0.00064; Games-Howell: *p*=0.001). By day 2, all conditions were significantly different (Welch’s ANOVA: *p*=0.0023; Games-Howell: EGF-control *p*=0.027, TGFβ1-control *p*=0.018, TGFβ1-EGF *p=*0.016) and the conditions were significantly different on day 3 as well (Welch’s Heteroscedastic F Test with Trimmed Means and Winsorized Variances: *p*=0.036; Games-Howell: EGF-control *p*=0.023, TGFβ1-control *p*=0.031, TGFβ1-EGF *p=*0.012). Cell viability was significantly different across all three groups (One-way ANOVA: *p*=1.78×10^−8^; Tukey Test: EGF-control *p*=1.05×10^−5^, TGFβ1-control *p*=4.6×10^−4^, TGFβ1-EGF *p*<1×10^−8^).

**Figure 3.**
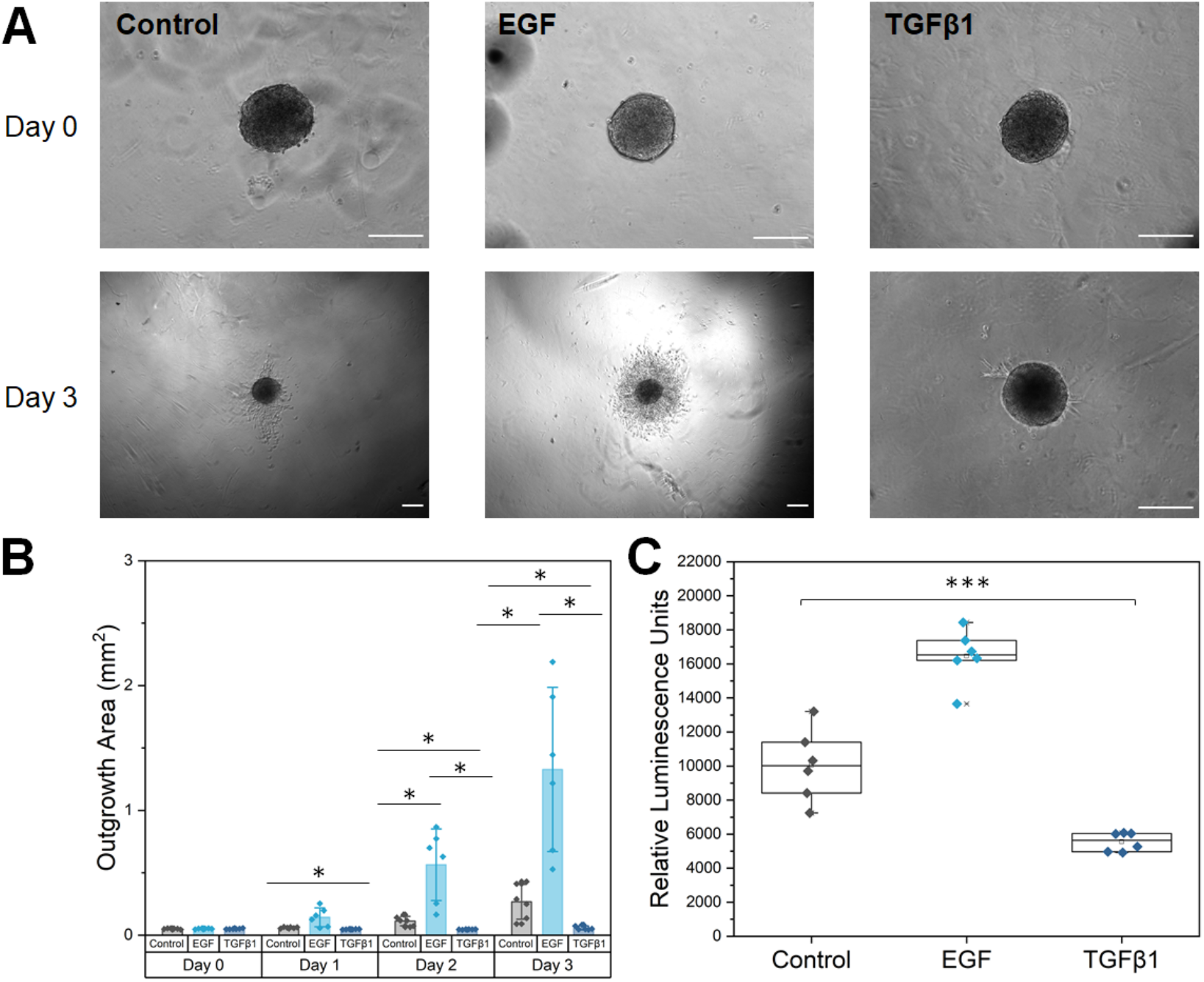
Epidermal growth factor (EGF) and transforming growth factor beta 1 (TGFβ1) modulate trophoblast outgrowth and viability. A) Representative phase images of HTR-8/SVneo trophoblast spheroid invasion at days 0 and 3. EGF and TGFβ1 were added to cell medium at day 0. Control samples contained no additional growth factors. B) Total spheroid outgrowth area at days 0-3 was quantified in Fiji and reported as averages for each condition. C) Cell viability data from CellTiter-Glo® 3D Cell Viability Assay measured in encapsulated spheroids on day 3. Data presented as mean ± standard deviation with individual data points shown. *: p<0.05. ***: p<0.001. n=6 samples per condition. Scale: 200 μm.

### 3.3. Cortisol has no effect on outgrowth area or viability

We subsequently used immunofluorescence staining of HTR-8/SVneo cells in two-dimensional cell culture well plates to validate their expression of glucocorticoid receptors. Notably, HTR-8/SVneo cells positively express glucocorticoid receptors (Fig. 4A). Further, we observed no evidence of non-specific antibody binding (samples stained without primary antibody) and observed only minimal background fluorescence. We subsequently quantified trophoblast spheroid outgrowth area and viability in the presence of cortisol. Although sample outgrowth area in the presence of cortisol for lower and physiological concentrations showed a decreasing trend in invasion by day 3 of culture, the effects were not significant. We observed no differences (*p*>0.05) in outgrowth area or viability (*p*=0.15) compared to control samples for lower (5 ng/mL and 20 ng/mL) concentrations at days 0 (*p*=0.88), 1 (*p*=0.23), 2 (*p*=0.08), and 3 (*p*=0.28) (Fig. 4B). Furthermore, for physiological (75 ng/mL and 150 ng/mL) concentrations of cortisol (Fig. 4C), we also observed no differences (*p*>0.05) in outgrowth area or viability at days 0 (*p*=0.52), 1 (*p*=0.83), 2 (*p*=0.06), and 3 (*p*=0.12) of culture.

**Figure 4.**
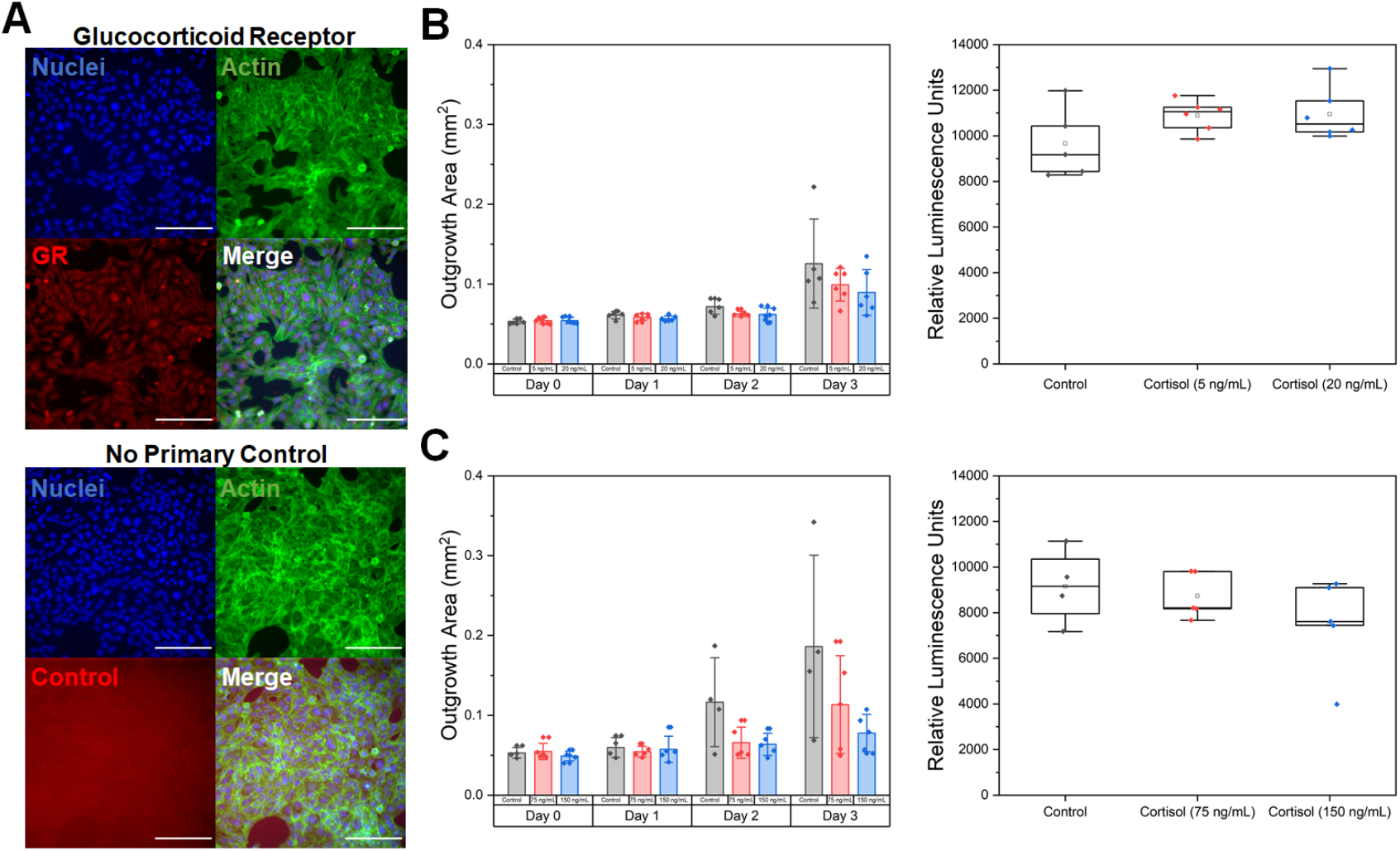
The effects of cortisol on trophoblast invasion. A) Representative fluorescent images of HTR-8/SVneo trophoblast cells cultured in 6-well plates. HTR-8/SVneo trophoblast cells express glucocorticoid receptors with cytoplasmic staining. The no primary control showed only minimal background fluorescence with no indication of non-specific binding. Green: Phalloidin (Actin). Red: Anti-Glucocorticoid Receptor. Blue: Hoechst (Nuclei). B) Total outgrowth area at days 0-3 and cell viability at day 3 for control, 5 ng/mL cortisol, or 20 ng/mL cortisol. C) Total outgrowth area at days 0-3 and cell viability at day 3 for control, 75 ng/mL cortisol, or 150 ng/mL cortisol. Data presented as mean ± standard deviation with individual data points shown. n=4-6 samples per condition. Scale: 200 μm.

## 4. Discussion

The objective of this study was to adapt a cell spheroid-based trophoblast invasion assay to probe how biomolecules from the maternal-fetal interface influence invasion. Here, we demonstrate that trophoblast invasion can be modulated through the use of known promoters and inhibitors of invasion. We then investigate cortisol, a stress hormone, and quantify invasion and cell viability in the presence of this molecule.

For these studies, we employ methacrylamide-functionalized gelatin (GelMA) hydrogels. Gelatin is denatured collagen and hence retains cell binding motifs and degradation sites that allow for cell attachment and matrix remodeling, key processes relevant to invasion (23, 27, 30–32). The endometrium contains collagens I, III, IV, V, and VI, rendering gelatin relevant in terms of its extracellular matrix composition (33, 34). Additionally, the biophysical properties of GelMA can be adjusted to fall within the regime of tissue stiffness. Recently, *Abbas et al.* used atomic force microscopy to demonstrate that the stiffness of the nonpregnant endometrium and placenta have an order of magnitude of 10^2^ Pa, whereas the decidua basalis, the region of the endometrium directly under the placenta that is invaded by trophoblast cells, exhibited an average stiffness on the order of 10^3^ Pa (29). Our hydrogel formulation falls within this range and replicates relevant mechanical properties of the tissue through which trophoblast cells will invade.

Unlike traditional two-dimensional wound healing assays or Boyden chamber assays, spheroid invasion assays in biomaterial platforms can replicate the three-dimensional structure of an invading blastocyst and provide a matrix through which cells can invade. We demonstrate reproducible spheroid formation using the HTR-8/SVneo cell line, resulting in spheroid size within the range of an invading blastocyst (28). We employ these trophoblast assays to interrogate differences in trophoblast outgrowth area and viability in the presence of biomolecules from the maternal-fetal interface. First, we selected known trophoblast invasion promoters and inhibitors: epidermal growth factor (EGF) and transforming growth factor β1 (TGFβ1) and observed quantifiable differences in trophoblast outgrowth area and cell viability. We show that soluble EGF stimulates trophoblast invasion and proliferation at concentrations used previously in literature (8, 14, 15). Additionally, soluble TGFβ1 inhibited trophoblast invasion and resulted in decreased proliferation, similar to what was reported previously in the literature (35, 36). Taken together, we show a multidimensional tissue engineering model can be used to quantify differences in invasion and proliferation using a small number of cells. Further, this platform is able to resolve effects of other biomolecules from the maternal-fetal interface known to modulate trophoblast invasion.

Subsequently, we included an investigation of the role of cortisol on trophoblast invasion. Stress is a physiological response caused by stressors and can be acute or chronic (20, 37). Significant stressors during pregnancy, including poverty, intimate partner violence, lack of social support, and racism, have been associated with increased risk for certain pregnancy disorders, including preterm birth, low birth weight, and preeclampsia (19–22). However, much remains unknown regarding how stress affects early events in pregnancy, such as trophoblast invasion. A benefit to our model system is that although the effects of stress on pregnancy outcomes may not be possible to study in humans or animals due to the complexity of the stress response, models of early implantation could allow us to study how stress affects trophoblast invasion using specific markers of stress (e.g., cortisol). Cortisol is a steroid hormone often used to assess the stress response and is functionally significant in processes of pregnancy (17, 18). When an individual experiences stressors, glucocorticoids are synthesized by the adrenal cortex in response to adrenocorticotrophic hormone (ACTH) from the pituitary gland (37). Although ACTH cannot cross the placenta, maternal glucocorticoids, including cortisol, can cross the placenta (38). Placental hydroxysteroid 11β dehydrogenase 2 (11β-HSD2) is an enzyme responsible for converting cortisol into its inactive form, cortisone, and protects the fetus from high maternal cortisol levels (39). However, increased maternal glucocorticoid levels can still lead to increased fetal exposure and may alter the activity of 11β-HSD2 as indicated by animal studies (38). Previous studies have shown that natural and synthetic glucocorticoids reduce trophoblast invasion and proliferation in two-dimensional cultures, wound healing assays, and Matrigel invasion assays (40–42).

We first investigated trophoblast invasion in the presence of cortisol at concentrations from *in vitro* studies (5-20 ng/mL) (43). Outgrowth area showed a decreasing trend in the presence of cortisol, although not statistically significant, with no changes in viability between groups. We next selected physiological concentrations of cortisol (75-150 ng/mL) comparable to what would occur during early pregnancy (18). We observed a similar trend, with no statistical differences in outgrowth area or viability despite physiologically relevant cortisol concentrations. To our knowledge, no studies to date have quantified cortisol levels in pregnant individuals with high stress levels; therefore, our selected concentrations may be lower than the required threshold to significantly reduce trophoblast invasion. Furthermore, 11β-HSD2 activity may have been able to convert these levels of cortisol into cortisone, which could explain why cortisol had a minimal effect on outgrowth area and viability (44). An investigation of 11β-HSD2 activity in the presence and absence of cortisol as well as a further exploration of cortisol dosages may provide additional insight. If no differences are found, it is possible that cortisol does not have an effect on trophoblast invasion or proliferation as previously found in the literature. Our system is three-dimensional and captures not only the spherical structure similar to an invading blastocyst but also allows cells to invade and remodel a surrounding gel matrix. Our system may be more applicable to what occurs physiologically and may not have as profound of an effect on invasion and proliferation as previously demonstrated in less complex models.

We recognize some limitations of our studies. These limitations provide exciting opportunities to improve our platforms to create models of increasing physiological relevance. We first acknowledge that the use of the HTR-8/SVneo cell line may contain a mixed population of cells and may not accurately mimic *in vivo* trophoblast behavior (45). For future investigations, the physiological relevance of our platform can be increased through the use of primary trophoblast cells. Secondly, we simplified an increased stress response to external stressors as equivalent to higher levels of cortisol. Although this strategy is easily testable in our platform, literature suggests that cortisol levels exhibit a dinural pattern and there is some individual variation in cortisol responsiveness based on mental health conditions, e.g., post-traumatic stress disorder (46–54). Future work on cortisol may be improved with temporal adjustments of cortisol concentrations to more appropriately mimic what occurs in the body.

## 5. Conclusions

Trophoblast invasion is a biological process essential to the establishment and success of a pregnancy. We employ GelMA hydrogels to perform advanced quantitative trophoblast invasion assays and demonstrate quantifiable differences in trophoblast outgrowth area and viability in the presence of known promoters and inhibitors of trophoblast invasion (EGF and TGFβ1). Further, we begin to probe how cortisol, a molecule from the maternal-fetal interface, may influence trophoblast invasion. The platform proposed herein will provide researchers with a unique tool critical for understanding implantation and will aid researchers in understanding mechanisms that dictate the success or failure of implantation.

## Acknowledgements

Research reported herein was supported by the National Institute of Diabetes and Digestive and Kidney Diseases of the National Institutes of Health under Award Numbers R01 DK099528 (B.A.C.H) and by the National Institute of Biomedical Imaging and Bioengineering of the National Institutes of Health under Award Numbers R21 EB018481 (B.A.C.H.). The content herein is solely the responsibility of the authors and does not necessarily represent the official views of the NIH. The authors also gratefully acknowledge additional funding provided by the Department of Chemical & Biomolecular Engineering and the Carl R. Woese Institute for Genomic Biology at the University of Illinois at Urbana-Champaign.

The authors would like to thank Dr. Jee-Wei Emily Chen (U. Illinois) for her incredible mentorship and assistance with method development and Dr. Sara Pedron (U. Illinois) for her assistance with NMR analysis. The authors would like to acknowledge the School of Chemical Sciences Cell Media Facility at the University of Illinois at Urbana-Champaign for assistance with cell media and the Core Facilities (Dr. Austin Cyphersmith) at the Institute for Genomic Biology at the University of Illinois at Urbana-Champaign for providing assistance with confocal imaging.

