## Supplemental Figure 1, Supplemental Figure 2 for "Tuning trophoblast invasion in a gelatin hydrogel via soluble cues from the maternal-fetal interface"

[1] Dept. of Bioengineering
[2] Dept. of Anthropology
[3] Beckman Institute for Advanced Science & Technology
[4] Dept. of Chemical and Biomolecular Engineering
[5] Carl R. Woese Institute for Genomic Biology
University of Illinois at Urbana-Champaign
Urbana, IL 61801

**Corresponding Author:**
B.A.C. Harley
Dept. of Chemical and Biomolecular Engineering
Carl R. Woese Institute for Genomic Biology
University of Illinois at Urbana-Champaign
110 Roger Adams Laboratory
600 S. Mathews Ave.
Urbana, IL 61801


**Additional Authors:**
K.B.H. Clancy
Dept. of Anthropology
University of Illinois at Urbana-Champaign
607 S. Mathews Ave.
Urbana, IL 61801


S.G. Zambuto
Dept. of Bioengineering
University of Illinois at Urbana-Champaign
1406 W. Green St
Urbana, IL 61801


***Supplemental 1. HTR-8/SVneo Cell Viability in Reduced Serum Conditions.***

HTR-8/SVneo cells were seeded onto 6-well plates at a density of 2 x $10^4$ cells/$cm^2$ and imaged every 24 hours in regular serum concentrations (5% fetal bovine serum (FBS)) or lower serum concentrations (2% FBS). After 72 hours, live/dead staining was performed to assess cell viability. We observed no obvious differences in cell morphology, growth, or viability between the two conditions.

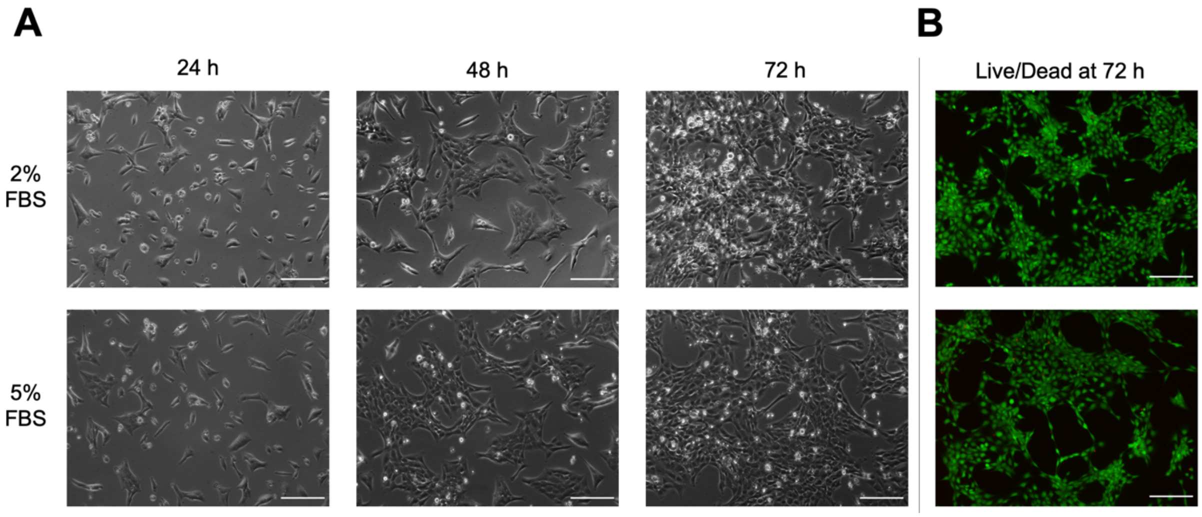


**Supplemental Figure 1.** Control experiment to verify cell HTR-8/SVneo growth and viability in reduced serum concentrations. A) Representative phase images of HTR-8/SVneo cells grown in well plates at 24, 48, and 72 hours after seeding cultured in growth medium supplemented with 2% or 5% fetal bovine serum (FBS). B) Live/dead images of HTR-8/SVneo cells at 72 hours after seeding. Scale: 200 μm.

***Supplemental 2. HTR-8/SVneo Trophoblast Spheroid Diameter.***

Encapsulated spheroids (n=18; 4,000 cells/spheroid) were measured on day 0 using Fiji. Spheroids were assumed to be spherical and diameter was quantified using the averages of 3 perimeter measurements and the equation for the circumference of a circle. Average spheroid diameter was found to be 270.3 ± 11.3 μm.

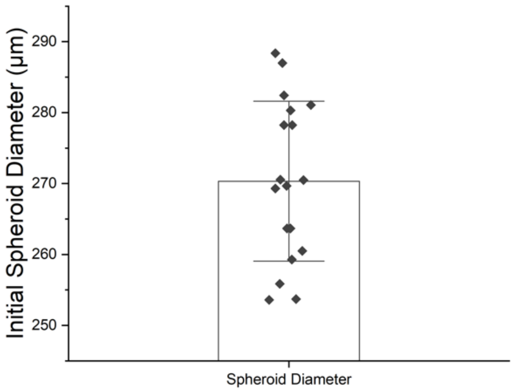


**Supplemental Figure 2.** Initial spheroid diameter on day 0 (encapsulation) for n=18 spheroids. Data presented as mean ± standard deviation with individual data points shown.
